# High seroprevalence of SARS-CoV-2 in white-tailed deer (*Odocoileus virginianus*) at one of three captive cervid facilities in Texas

**DOI:** 10.1101/2022.01.05.475172

**Authors:** Christopher M. Roundy, Chase M. Nunez, Logan F. Thomas, Lisa D. Auckland, Wendy Tang, Jack J. Richison, Breanna R. Green, Clayton D. Hilton, Michael J. Cherry, Alex Pauvolid-Correa, Gabriel L. Hamer, Walter E. Cook, Sarah A. Hamer

## Abstract

Free-ranging white-tailed deer (*Odocoileus virginanus*) across the United States are increasingly recognized as involved in SARS-CoV-2 transmission cycles. Through a cross-sectional study of 80 deer at three captive cervid facilities in central and southern Texas, we provide evidence of 34 of 36 (94.4%) white-tailed deer at a single captive cervid facility seropositive for SARS-CoV-2 by neutralization assay (PRNT_90_), with endpoint titers as high as 1280. In contrast, all tested white-tailed deer and axis deer (*Axis axis*) at two other captive cervid facilities were seronegative, and SARS-CoV-2 RNA was not detected in respiratory swabs from deer at any of the three facilities. These data support transmission among captive deer that cannot be explained by human contact for each infected animal, as only a subset of the seropositive does had direct human contact. The facility seroprevalence was more than double of that reported from wild deer, suggesting that the confined environment may facilitate transmission. Further exploration of captive cervids and other managed animals for their role in the epizootiology of SARS-CoV-2 is critical for understanding impacts on animal health and the potential for spillback transmission to humans or other animal taxa.

**Importance:** As SARS-CoV-2 vaccine coverge of the human population increases and variants of concern continue to emerge, identification of the epidemiologic importance of animal virus reservoirs is critical. We found that nearly all (94.4%) of the captive white-tailed deer at a cervid facility in central Texas had neutralizing antibodies for SARS-CoV-2. This seroprevalence is over double than that which has been reported from free-ranging deer from other regions of the US. Horizontal transmission among deer may be facilitated in confinement. Tracking new infections among wild and confined deer is critical for understanding the importance of animal reservoirs for both veterinary and human health.

## Introduction

Despite many public health preventative measures and the advance of vaccination worldwide, epidemic waves of severe acute respiratory syndrome coronavirus 2 (SARS-CoV-2) caused by more transmissible and antibody evasive genetic variants of concern have sustained the ongoing virus circulation (1, 2). The susceptibility to infection and potential capacity to transmit SARS-CoV-2 in domestic and wildlife species have been increasingly reported (3-6); this understanding is especially critical given wild animals remain unvaccinated and could serve as viral reservoirs.

Recent experimental studies have demonstrated that the white-tailed deer (*Odocoileus virginianus*) is highly susceptible to infection and can transmit SARS-CoV-2 by contact and vertical transmission (7-11). Notably, the animals that were infected by contact shed infectious virus in nasal secretions for up to seven days, indicating that SARS-CoV-2 can be propagated for prolonged periods in a herd after a single infection (8). Approximately 37-40% seroprevalence of wild white-tailed deer to SARS-CoV-2 was reported across three Midwestern states (9) and Texas (12). Additionally, one third of 283 free-living and captive white-tailed deer from Iowa in the post-pandemic period had SARS-CoV-2 RNA in their lymph nodes including several lineages circulating in humans, suggesting multiple spillover events followed by deer-to-deer transmission (13), and over one third (35.8%) of 360 wild deer from Ohio had SARS-CoV-2 positive nasal swabs comprised of several variants, including those that were domininat as well as uncommon in the human population (14).

The captive cervid industry arose from growing demand for deer and deer products and involves the raising, or containment propogation of native and non-native cervids in privately-maintained facilities, often involving close human-deer contacts. The operations of captive cervid facilities are diverse and include venison and antler production, sales to other cervid operations, coordinated and uncoordinated propogation, providing for commercial fenced hunting preserves, genetic enhancement of private deer herds, and more. There are approximately 10,000 deer breeding and/or hunting facilities in North America (15), most of which include white-tailed deer. The industry is particularly important in Texas where the deer hunting and breeding industry generates over $1.6 billion in economic activity annually (16). Given the recent findings of wild white-tailed deer infection with SARS-CoV-2, combined with the precedent from mink that captive animal facilities may be high risk transmission environments (17-19), our objective was to investigate the SARS-CoV-2 exposure and infection status of captive cervids in Texas.

## Materials & Methods

*Captive cervid facilities*. Between September-October 2021, three captive cervid facilities in Texas were sampled. Facility A is a private white-tailed deer breeding and hunting ranch in Guadalupe Co., TX, with approximately 207 captive white-tailed deer (57 bucks, 150 does) used for breeding and research purposes; twenty-one of these does were previously enrolled in experimental anthrax vaccine work that involved being handled individually for <1 minute for blood collection at approximately monthly time points from November 2020 – July 2021, with each hands-on interaction lasting less than one minute. The facility includes thirteen pens ranging from 0.4–1.6 acres; twelve for housing breed stock and one for housing research animals, all of which have fenceline contact with two to five occupied deer pens. Once-a-day scheduled feedings in each pen were provided by either the ranch foreman or owner. Facility B is a private 7.2-acre pen in Montgomery Co., TX, housing approximately 22 Axis deer (*Axis axis*; 9 bucks, 13 does) and 7 fallow deer (*Dama dama*; 2 bucks, 5 does) for breeding purposes. The exotic cervids’ ages ranged from 0.5 – 7.5 years. Human contact consisted of approximately once a week feeder checks by one of five individuals. Facility C contains a reserch deer herd owned by Texas A&M University-Kingsville in Kleberg Co, TX, with approximately 40 white-tailed deer (19 bucks, 21 does) ranging in age from 0.5–13.5 years used for teaching and research. The facility consists of six deer pens ranging from 0.3–1.1 acres. Approximately 25 of the deer from Facility C had been previously enrolled in the same anthrax vaccine research as Facility A and all except the eight fawns from 2021 have previously been involved in ecology and management studies.

*Sample collection*. During the course of regular veterinary care of deer under physical restraint or chemical immobilization, samples were collected for the SARS-CoV-2 investigation. Two swabs were collected from deer using polyester-tipped applicators with polystyrene handles (Puritan Medical Products, Guilford, ME, USA) and stored in viral transport media (VTM; made following CDC SOP#: DSR-052-02): (i) respiratory (which consisted of one nasal swab and one oral swab placed together into the vial) and (ii) rectal. Blood samples were collected using jugular venipuncture into no-additive or clot activator tubes. All samples were collected in accordance with the relevant guidelines and regulations approved by the TAMU’s Institutional Animal Care and Use Committee and Clinical Research Review Committee (2018-0460 CA).

*Molecular diagnostics*. Aliquots of VTM supernatant were subjected to RNA/DNA extraction by MagMAX™ CORE Nucleic Acid Purification Kit (ThermoFisher Scientific, Waltham, MA). An aliquot of purified nucleic acid was tested by SARS-CoV-2 specific real time qRT-PCR to amplify the RdRp gene (20).

*Plaque reduction neutralization tests* (PRNT). Serum samples were tested by PRNT to quantify antibodies able to neutralize the formation of SARS-CoV-2 plaques on Vero cell cultures following standard protocols (21) in a BSL3 laboratory. Serum samples were heat inactivated and screened at a dilution of 1:10 and those that neutralized SARS-CoV-2 viral plaques by at least 90%, when compared to the virus control, were further tested at serial two-fold dilutions starting at 1:10 to 1:320 to determine 90% endpoint titers. The SARS-CoV-2 Isolate USAIL1/2020, NR 52381 was used to prepare virus stocks (BEI Resources, Manassas, VA).

## Results

Between September 15, 2021 and November 29, 2021, we collected and tested respiratory and rectal swabs and serum of 80 deer from 3 captive cervid facilities in central and southern Texas. Across all 80 deer from all 3 facilities, respiratory and rectal swabs were RT-PCR negative. At Facilty A, 34 of 36 (94.4%) of adult deer (mean age of 3.5 years among bucks, 4.5 years for does) were seropositive for SARS-CoV-2 with endpoint titers as high as 1280 (Table 1); a subset of the does only had direct human contact. At Facilities B and C, all deer were seronegative.

**Table 1.** Diagnostic details of captive cervids tested for SARS-CoV-2 RNA and antibodies at three facilities in central and South Texas, Sept-Oct, 2021.

| Site | County | Species | Sex | No. tested | RT-qPCR positives |  | PRNT90 |  |  |
| --- | --- | --- | --- | --- | --- | --- | --- | --- | --- |
|  |  |  |  |  | Respiratory swab | Rectal swab | Seropositives (%) | Titer range | Titer geometric mean |
| Ranch A | Guadalupe, TX | *Odocoileus virginianus* | F | 21 | 0 | 0 | 19 (90.50%) | 20 - 320 | 69.13 |
|  |  |  | M | 15 | 0 | 0 | 15 (100%) | 20 - 1,280 | 152.77 |
| Ranch B | Montgomery, TX | *Axis axis* | F | 8 | 0 | 0 | 0 (0%) | <10 | <10 |
|  |  |  | M | 7 | 0 | 0 | 0 (0%) | <10 | <10 |
|  |  | *Dama dama* | M | 1 | 0 | 0 | 0 (0%) | <10 | <10 |
| Ranch C | Kleberg, TX | *Odocoileus virginianus* | F | 14 | 0 | 0 | 0 (0%) | <10 | <10 |
|  |  |  | M | 15 | 0 | 0 | 0 (0%) | <10 | <10 |

## Discussion

We found that nearly all (94.4%) the tested white-tailed deer at a single captive cervid facility in Texas were positive for SARS-CoV-2 neutralizing antibodies, yet they were negative for viral RNA in respiratory and rectal swabs. In contrast, all the tested deer at two other captive cervid facilities were seronegative and negative for viral RNA. Although clinical data were not systematically recorded for research purposes, the seropositive deer were observed at least once daily and were not reported to show any clinical signs of disease at any time prior to, during, or after they were sampled.

Many, but not all, of the seropositive animals were used previously in research to evaluate an experimental anthrax vaccine. Nineteen of the twenty-one white-tailed does tested at Facility A were either given experimental vaccine or the commercially available Sterne spore vaccine, while all of the fifteen white-tailed bucks tested were neither researched or handled by humans prior to sampling. Furthermore, many of the seronegative individuals from Facility C were enrolled in the same anthrax vaccine immunogeneticity study and showed no titers to SARS-CoV-2, indicating that cross-reactivity from the various anthrax vaccines is unlikely.

At Facilty A the ranch manager and owner both disclosed that they tested positive for SARS-CoV-2 on 8/4/21 and 8/6/21, respectively, approximately 5 weeks prior to the collection of samples from the deer. At Facility C, at least one animal caretaker was tested positive for SARS-CoV-2 on 7/9/21, approximately 16 weeks prior to the sampling. No previously confirmed cases of SARS-CoV-2 occured in the six humans that manage the deer at Facility B.

The level of exposure among deer at the captive cervid facility (94.4%) is more than double that which has recently been reported across several studies of free-ranging individuals (9, 12-14); one explanation may be that onward transmission among deer is facilitated by the confined environment. The circulation of SARS-CoV-2 among captive animals could result in eventual spillover to other wildlife species with unknown impact for the conservation and public health, considering the high rates of mutation and recombination observed among coronaviruses (22). A One Health approach is critical to advance the understanding of the potential impact of human-animal interface for SARS-CoV-2 transmission, maintenance and evolution.

## Acknowledgements

AgriLife Research provided funding. The following reagent was deposited by the Centers for Disease Control and Prevention and obtained through BEI Resources, NIAID, NIH: SARS-Related Coronavirus 2, Isolate USAIL1/2020, NR 52381.

## References

1. Galloway SE, Paul P, MacCannell DR, Johansson MA, Brooks JT, MacNeil A, Slayton RB, Tong S, Silk BJ, Armstrong GL, Biggerstaff M, Dugan VG. 2021. Emergence of SARS-CoV-2 B.1.1.7 lineage - United States, December 29, 2020-January 12, 2021. MMWR Morb Mortal Wkly Rep 70:95–99.

2. Dejnirattisai W, Zhou D, Supasa P, Liu C, Mentzer AJ, Ginn HM, Zhao Y, Duyvesteyn HME, Tuekprakhon A, Nutalai R, Wang B, Lopez-Camacho C, Slon-Campos J, Walter TS, Skelly D, Costa Clemens SA, Naveca FG, Nascimento V, Nascimento F, Fernandes da Costa C, Resende PC, Pauvolid-Correa A, Siqueira MM, Dold C, Levin R, Dong T, Pollard AJ, Knight JC, Crook D, Lambe T, Clutterbuck E, Bibi S, Flaxman A, Bittaye M, Belij-Rammerstorfer S, Gilbert SC, Carroll MW, Klenerman P, Barnes E, Dunachie SJ, Paterson NG, Williams MA, Hall DR, Hulswit RJG, Bowden TA, Fry EE, Mongkolsapaya J, Ren J, Stuart DI, Screaton GR. 2021. Antibody evasion by the P.1 strain of SARS-CoV-2. Cell 184:2939–2954 e9.

3. Randazzo W, Cuevas-Ferrando E, Sanjuan R, Domingo-Calap P, Sanchez G. 2020. Metropolitan wastewater analysis for COVID-19 epidemiological surveillance. Int J Hyg Environ Health 230:113621.

4. Bosco-Lauth AM, Root JJ, Porter SM, Walker AE, Guilbert L, Hawvermale D, Pepper A, Maison RM, Hartwig AE, Gordy P, Bielefeldt-Ohmann H, Bowen RA. 2021. Peridomestic Mammal Susceptibility to Severe Acute Respiratory Syndrome Coronavirus 2 Infection. Emerg Infect Dis 27:2073–2080.

5. Hamer SA, Pauvolid-Correa A, Zecca IB, Davila E, Auckland LD, Roundy CM, Tang W, Torchetti MK, Killian ML, Jenkins-Moore M, Mozingo K, Akpalu Y, Ghai RR, Spengler JR, Barton Behravesh C, Fischer RSB, Hamer GL. 2021. SARS-CoV-2 Infections and Viral Isolations among Serially Tested Cats and Dogs in Households with Infected Owners in Texas, USA. Viruses 13:938.

6. Karikalan M, Chander V, Mahajan S, Deol P, Agrawal RK, Nandi S, Rai SK, Mathur A, Pawde A, Singh KP, Sharma GK. 2021. Natural infection of Delta mutant of SARS-CoV-2 in Asiatic lions of India. Transbound Emerg Dis doi:10.1111/tbed.14290.

7. Cool K, Gaudreault NN, Morozov I, Trujillo JD, Meekins DA, McDowell C, Carossino M, Bold D, Kwon T, Balaraman V, Madden DW, Artiaga BL, Pogranichniy RM, Sosa GR, Henningson J, Wilson WC, Balasuriya UBR, García-Sastre A, Richt JA. 2021. Infection and transmission of SARS-CoV-2 and its alpha variant in pregnant white-tailed deer. bioRxiv doi:10.1101/2021.08.15.456341.

8. Palmer MV, Martins M, Falkenberg S, Buckley A, Caserta LC, Mitchell PK, Cassmann ED, Rollins A, Zylich NC, Renshaw RW, Guarino C, Wagner B, Lager K, Diel DG. 2021. Susceptibility of white-tailed deer (Odocoileus virginianus) to SARS-CoV-2. J Virol 95.

9. Chandler JC, Bevins SN, Ellis JW, Linder TJ, Tell RM, Jenkins-Moore M, Root JJ, Lenoch JB, Robbe-Austerman S, DeLiberto TJ, Gidlewski T, Kim Torchetti M, Shriner SA. 2021. SARS-CoV-2 exposure in wild white-tailed deer (Odocoileus virginianus). Proc Natl Acad Sci U S A 118.

10. Martins M, Boggiatto PM, Buckley A, Cassmann ED, Falkenberg S, Caserta LC, Fernandes MHV, Kanipe C, Lager K, Palmer MV, Diel DG. 2021. From Deer-to-Deer: SARS-CoV-2 is efficiently transmitted and presents broad tissue tropism and replication sites in white-tailed deer. bioRxiv doi:10.1101/2021.12.14.472547:2021.12.14.472547.

11. Sit THC, Brackman CJ, Ip SM, Tam KWS, Law PYT, To EMW, Yu VYT, Sims LD, Tsang DNC, Chu DKW, Perera R, Poon LLM, Peiris M. 2020. Infection of dogs with SARS-CoV-2. Nature doi:10.1038/s41586-020-2334-5.

12. Palermo PM, Orbegozo J, Watts DM, Morrill JC. 2021. SARS-CoV-2 Neutralizing Antibodies in White-Tailed Deer from Texas. Vector Borne Zoonotic Dis doi:10.1089/vbz.2021.0094.

13. Kuchipudi SV, Surendran-Nair M, Ruden RM, Yon M, Nissly RH, Nelli RK, Li L, Jayarao BM, Vandegrift KJ, Maranas CD, Levine N, Willgert K, Conlan AJK, Olsen RJ, Davis JJ, Musser JM, Hudson PJ, Kapur V. 2021. Multiple spillovers and onward transmission of SARS-CoV-2 in free-living and captive white-tailed deer. bioRxiv doi:10.1101/2021.10.31.466677:2021.10.31.466677.

14. Hale VL, Dennis PM, McBride DS, Nolting JM, Madden C, Huey D, Ehrlich M, Grieser J, Winston J, Lombardi D, Gibson S, Saif L, Killian ML, Lantz K, Tell R, Torchetti M, Robbe-Austerman S, Nelson MI, Faith SA, Bowman AS. 2021. SARS-CoV-2 infection in free-ranging white-tailed deer. Nature doi:10.1038/s41586-021-04353-x.

15. Anderson DP, B. J. Frosch, and J. L. Outlaw. 2007. Economic impact of the Texas deer breeding industry. Texas A&M University, Agricultural and Food Policy Center Research Report 07-3, College Station, USA.,

16. Outlaw JL, Anderson DP, Earle ML, Richardson JW. 2017. Economic Impact of the Texas Deer Breeding and Hunting Operations. Texas A&M University, College Station, TX.

17. Rasmussen TB, Fonager J, Jorgensen CS, Lassauniere R, Hammer AS, Quaade ML, Boklund A, Lohse L, Strandbygaard B, Rasmussen M, Michaelsen TY, Mortensen S, Fomsgaard A, Belsham GJ, Botner A. 2021. Infection, recovery and re-infection of farmed mink with SARS-CoV-2. PLoS Pathog 17:e1010068.

18. Lu L, Sikkema RS, Velkers FC, Nieuwenhuijse DF, Fischer EAJ, Meijer PA, Bouwmeester-Vincken N, Rietveld A, Wegdam-Blans MCA, Tolsma P, Koppelman M, Smit LAM, Hakze-van der Honing RW, van der Poel WHM, van der Spek AN, Spierenburg MAH, Molenaar RJ, de Rond J, Augustijn M, Woolhouse M, Stegeman JA, Lycett S, Munnink BBO, Koopmans MPG. 2021. Adaptation, spread and transmission of SARS-CoV-2 in farmed minks and associated humans in the Netherlands. Nature Communications 12:6802.

19. Eckstrand CD, Baldwin TJ, Rood KA, Clayton MJ, Lott JK, Wolking RM, Bradway DS, Baszler T. 2021. An outbreak of SARS-CoV-2 with high mortality in mink (Neovison vison) on multiple Utah farms. PLoS Pathog 17:e1009952.

20. Corman VM, Landt O, Kaiser M, Molenkamp R, Meijer A, Chu DK, Bleicker T, Brunink S, Schneider J, Schmidt ML, Mulders DG, Haagmans BL, van der Veer B, van den Brink S, Wijsman L, Goderski G, Romette JL, Ellis J, Zambon M, Peiris M, Goossens H, Reusken C, Koopmans MP, Drosten C. 2020. Detection of 2019 novel coronavirus (2019-nCoV) by real-time RT-PCR. Euro Surveill 25.

21. Beaty BC, C.H.; Shope, R.E. 1995. Arboviruses, p 189–212. In Lennette EH, Schmidt, N.J. (ed), Viral, Rickettsial, and Chlamydial Infections. American Public Health Association, Washington, DC, USA.

22. Brian DA, Baric RS. 2005. Coronavirus genome structure and replication. Curr Top Microbiol Immunol 287:1–30.

